# Combination therapy of metformin with sodium selenite reverses effects of monotherapy on nitric oxide production, IL-1β and TNF-α release, and upregulates relative expression of Bcl-2 by LPS-activated human primary monocytes in T-cell acute lymphoblastic leukemia

**DOI:** 10.1101/2022.01.04.475000

**Authors:** Fella Rostane, Nidel Sari, Ilyes Bali, Rabia Messali, Zeyneb Hadjidj, Maroua Miliani, Imène Belhassena, Charazed El Mezouar, Mourad Aribi

## Abstract

**Objectives:** We examined the influence of the *ex vivo* combination therapy of metformin (Met, 1,1-dimethylbiguanide hydrochloride) with sodium selenite (Ss, Na^2^SeO^3^) on the changes in the production of nitric oxide (NO) and selected cytokines by circulating monocytes (MOs) during T-cell acute lymphoblastic leukemia (T-ALL).

**Methods:** Assays were performed on MO cell samples isolated from children with T-ALL.

**Results:** Met+Ss combination therapy reversed the Ss effect on the upregulation of NO production. Both Met+Ss and Ss treatment alone induced a significant downregulation of extracellular calcium ions consumption (_ec_Ca^2+^) levels. Additionally, Met treatment induced a significant upregulation of IL-1β and TNF-α production; such effects were significantly reversed after combination with Ss treatment. Moreover, Met+Ss induced no significant effect on the production of IL-10, IL-6 and TNF-α, but a slight increase in IFN-γ levels. Furthermore, treatment with Ss alone induced a slight increase of IFN-γ. Finally, Met+Ss induced a marked upregulation of relative Bcl-2 expression in MOs.

**Conclusions:** Met+Ss combination therapy results in downregulation of NO production, IL-1β and TNF-α release as well as in upregulation of the relative expression levels of Bcl-2-associated survival of primary MOs in human T-ALL.

## 1. Introduction

T-ALL is an aggressive hematological tumor, characterized by the clonal proliferation of immature T lymphoid line hematopoietic precursor that arise from the thymus and infiltrate into the bone marrow and peripheral blood [1]. The T-cell malignant transformation results from a multistep oncogenic process which affects the normal mechanisms that control cell growth, proliferation, survival, and differentiation during thymocyte development [2]. Additionally, it has been reported that the development and progression of malignant cells can be affected by changes in the levels of NO [3] and cytokines profile [4] derived from host cells, including immune cells, like MOs.

More recently, MOs have emerged as the main cells involved in cancer immunopathogenesis [5], and can exhibit a dual role and contributes either to pro or antitumoral immunity [6]. Their functional activities, including their ability to produce NO and cytokines, have been observed to tend to change during the course of various treatment [7]. Of note, Met, a biguanide class drug known by its anti-hyperglycemic action [8], has been highlighted for it notably anti-inflammatory and preventive effect against various malignant tumors [9]. It seems to have pleiotropic properties, by exerting an anti-tumor effect either indirectly, by interfering with the insulin signaling pathways, or directly on tumor cells, by various signaling pathways [9,10]. Therefore, know of its effects on immune cells can undoubtedly help to develop new therapeutic approaches.

Similarly to Met, a particular therapeutic interest has been attributed to selenium, an essential micronutrient, for its pleiotropic and immunomodulatory effects [11,12], as well as for its potent role in inhibition of expression of LPS induced proinflammatory cytokines [13]. It also enhances immunity and the function of cytotoxic effector cells and the production of antibodies [14,15]. Additionally, it has been shown that selenium treatment suppresses leukemia by triggering apoptosis of primary human and murine cancer stem-like cells (CSC) through arachidonic acid metabolism and the subsequent action of endogenous eicosanoids [16]. Moreover, Ss supplementation can influence Met effects in normal conditions [8].

Of note, it has been reported that combination therapy has been considered the cornerstone of cancer therapy, promoting improved therapeutic efficacy compared to the monotherapy approach, as it targets key pathways in a synergistic or additive manner [17]. Based on the aforementioned, we examined in this first report the influence of the *ex vivo* combination therapy of Met with selenium, in the form of Ss (Ss, Na_2_SeO_3_), on changes in the production of NO and selected cytokines by circulating MOs during T-ALL.

## 2. Materials and methods

### 2.1 Study design and cell samples

We studied the *ex vivo* effect of Met, combined or not with Ss, on the modulation of the production of NO and selected cytokines by primary MOs isolated from peripheral blood mononuclear cells (PBMCs) of volunteer children suffering from T-ALL admitted at the Pediatric Department of Tlemcen Specialized Mother & Child Hospital Establishment *(Etablissement Hospitalier Spécialisé [EHS] Mère-Enfant).* Written informed consent was obtained from the parents and the study was approved by the Ethics Committee of Tlemcen University. MOs samples were devised into four groups, including untreated cell (UNT) controls, and cells treated with Met alone or Ss alone or Met combined with Ss (Met+Ss). NO production, _ec_Ca^2+^ levels and the release of pro-and anti-inflammatory cytokines were examined in MOs supernatant cultures. Bcl-2 levels were measured in intracellular MOs. Experiments were repeated at least three times. Rationale and flow-chart of the current study are shown in Figure 1.

**Figure 1.**
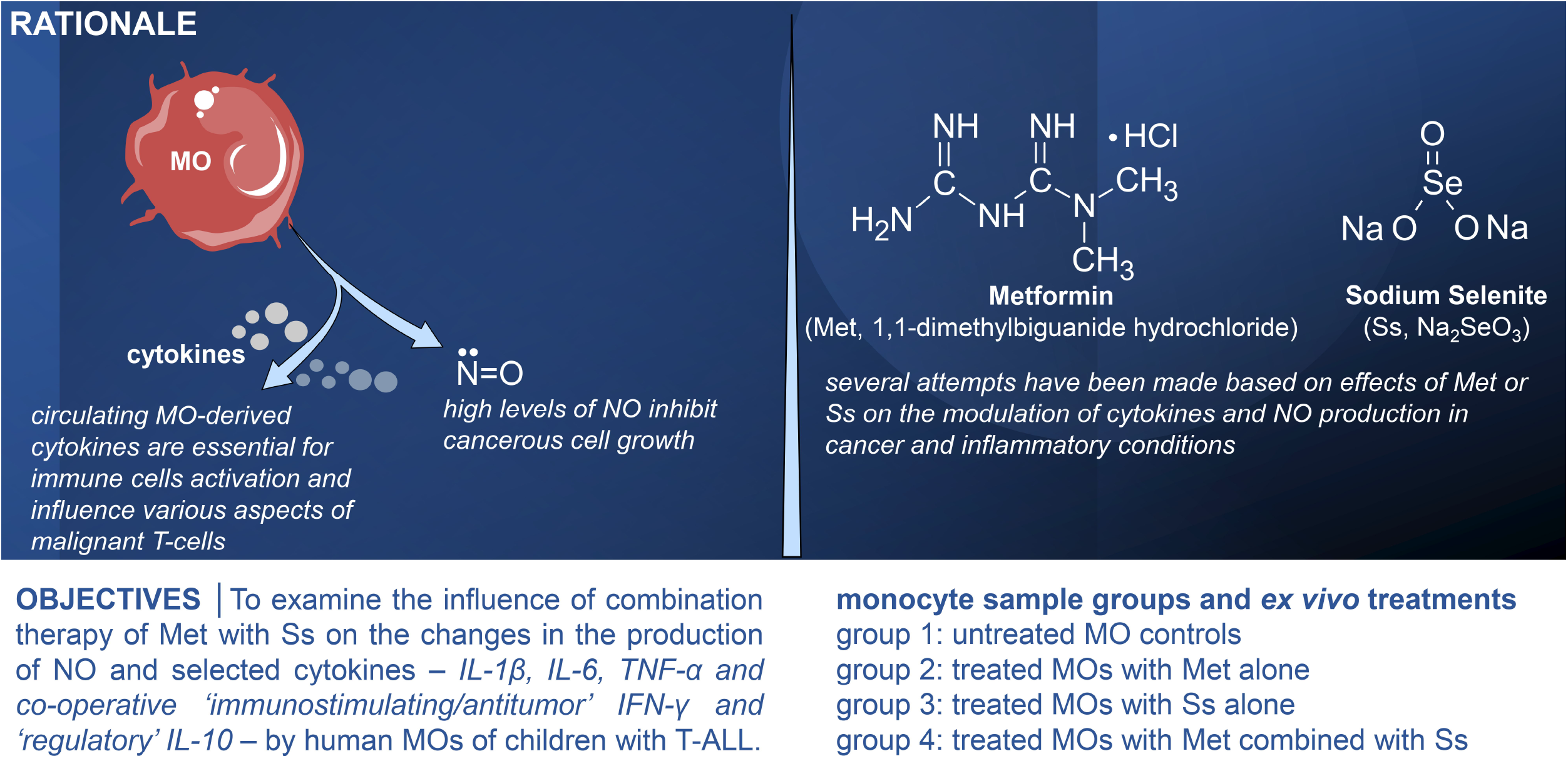
Rationale and flow-chart of the current study. The current study was focused on the determination of the *ex vivo* effects of combination therapy of metformin with sodium selenite on functional activities of human primary MOs isolated from volunteer children with T-ALL, including the production levels of NO and selected cytokines, _ec_Ca^2+^ levels, as well as the MO Bcl-2-associated survival expression levels. The study was conducted in four groups of MO cell samples, including untreated cell controls (UNT) and cells treated by metformin alone (Met) or sodium selenite alone (Ss) or Met combined with Ss (Met+Ss). Bcl-2: B cell lymphoma-2, _ec_Ca^2+^: extracellular calcium ion consumption, Met: metformin, MO: monocyte, NO: nitric oxide, Ss: sodium selenite, T-ALL: T-cell acute lymphoblastic leukemia. Cell illustrations and arrows were freely provided from Servier Medical Art (https://smart.servier.com/).

### 2.2 Cell isolation and purification, and MOs treatment

PBMCs were purified by Ficoll-Histopaque density gradient centrifugation (Ficoll-Histopaque 1077 Sigma-Aldrich, St Louis, MO, USA)] and cultured at 37 °C and 5% CO_2_ with humid atmosphere in minimum essential medium (MEM), containing 5% FBS, 50 μg/mL penicillin, 50 μg/mL streptomycin, 2 mM L-glutamine [18]. MOs were isolated from PBMCs by plastic adherence method [19]. The non-adherent cells were removed and adherent MOs subjected or not to treatment. Cell viability was evaluated by using the Trypan Blue Exclusion Test (TBET) [20], and the cell suspensions purity was higher than 90% as assessed by fluorescent staining with PhycoErytherin (PE)-anti-human CD14 antibody (BD Biosciences, San Diego, CA, USA) using a Floid Cell Imaging Station (Thermo Fisher Scientific, MA USA) [21,22]. 2 × 10^5^ MOs were stimulated for 24 h at 37 °C and 5% CO_2_ with 1 μg/mL LPS (*Escherichia coli* 0127:B8, Sigma-Aldrich). After LPS elimination, MOs were left either untreated (UNT) or treated with a dose of 1 μM Met or with 5 ng/mL (≅30 nmol/L Ss) Ss in a final volume of 200 μL MEM, or with Met combined with Ss (Met+Ss), while respecting the same concentrations of each in the final volume of culture medium [8].

### 2.3 NO assay

The levels of NO production were determined by measuring the physiologically stable oxidative metabolite (NOx, nitrite [NO^2^-] and nitrate [NO^3^-]) accumulation by the sensitive Griess method, as described [23,24]. Absorbance was spectrophotometrically determined at 540 nm on the ELISA plate reader (Biochrom Anthos 2020, Cambridge, UK).

### 2.4 _ec_Ca^2+^assay

The _ec_Ca^2+^ concentrations were measured spectrophotometrically using calcium colorimetric assay kit from Sigma-Aldrich (Sigma-Aldrich, MAK022-1KT). _ec_Ca^2+^ levels were determined as follows: _ec_Ca^2+^ = [Ca^2+^]_CM_ - [Ca^2+^]_MO_, where [Ca^2+^]_CM_ correspond to the concentration of calcium ions in culture medium alone, and [Ca^2+^]_MO_ correspond to the concentration of calcium ions in MO culture supernatants. _ec_Ca^2+^levels were expressed as μmol/2 × 10^5^ cells/mL.

### 2.5 Bcl-2 expression assay

MOs were washed with phosphate-buffered-saline (PBS), fixed and permeabilized with PBS containing 4% paraformaldehyde and 0.5% Triton X-100. The well plates were washed with 0.05% PBS-Tween 20 solution (v/v), and then Bcl-2 primary detector antibodies were added to each well, and incubated for 2 h at room temperature. Thereafter, horseradish peroxidase (HRP)-conjugated secondary antibody was added. After 1 h incubation, 3,3’,5,5’-tétraméthylbenzidine (TMB) was added. Finally, a last incubation was made for 30 min in the dark. The chromogenic reaction was stopped by the addition of 50 μL 2 N phosphoric acid solution. The absorbance was read immediately at 450 nm using ELISA plate reader (Biochrom Anthos 2020, Cambridge, UK) [25].

### 2.6 Cytokine assays

The concentration of IFN-γ, IL1-β, IL-6, IL-10 and TNF-α were determined in 24 h culture supernatant of UNT control and treated MOs using respective ELISA kits (Sigma-Aldrich kit, USA), according to the manufacturer’s instructions. Absorbance was measured at 450 nm in triplicate, by ELISA plate reader (Biochrom Anthos 2020, Cambridge, UK).

### 2.7 Statistical analyses

Comparisons were performed by one-way analysis of variance (ANOVA), followed by Tukey’s multiple comparisons test. *P*-values less than 0.05 were considered significant. Statistical analyses and Figures conception were carried out using GraphPad Prism software version 8.0.1 (San Diego, CA, USA).

## 3. Results

### 3.1 Met+Ss combination therapy reverses the Ss effect on MO NO production

As depicted in Figure 2, we observe that Met treatment has no significant effect on NO production by MOs as compared to Met-untreated MOs *(p* > 0.05). However, when combined to Ss, Met induced a markedly downregulation of NO production *(p* = 0.0027 by Tukey’s multiple comparisons test). Conversely, Ss induced upregulation of NO production by MOs from T-ALL patients, when used alone *(p* = 0.0015 by Tukey’s multiple comparisons test). *P*-value was < 0.001 by one-way ANOVA test.

**Figure 2.**
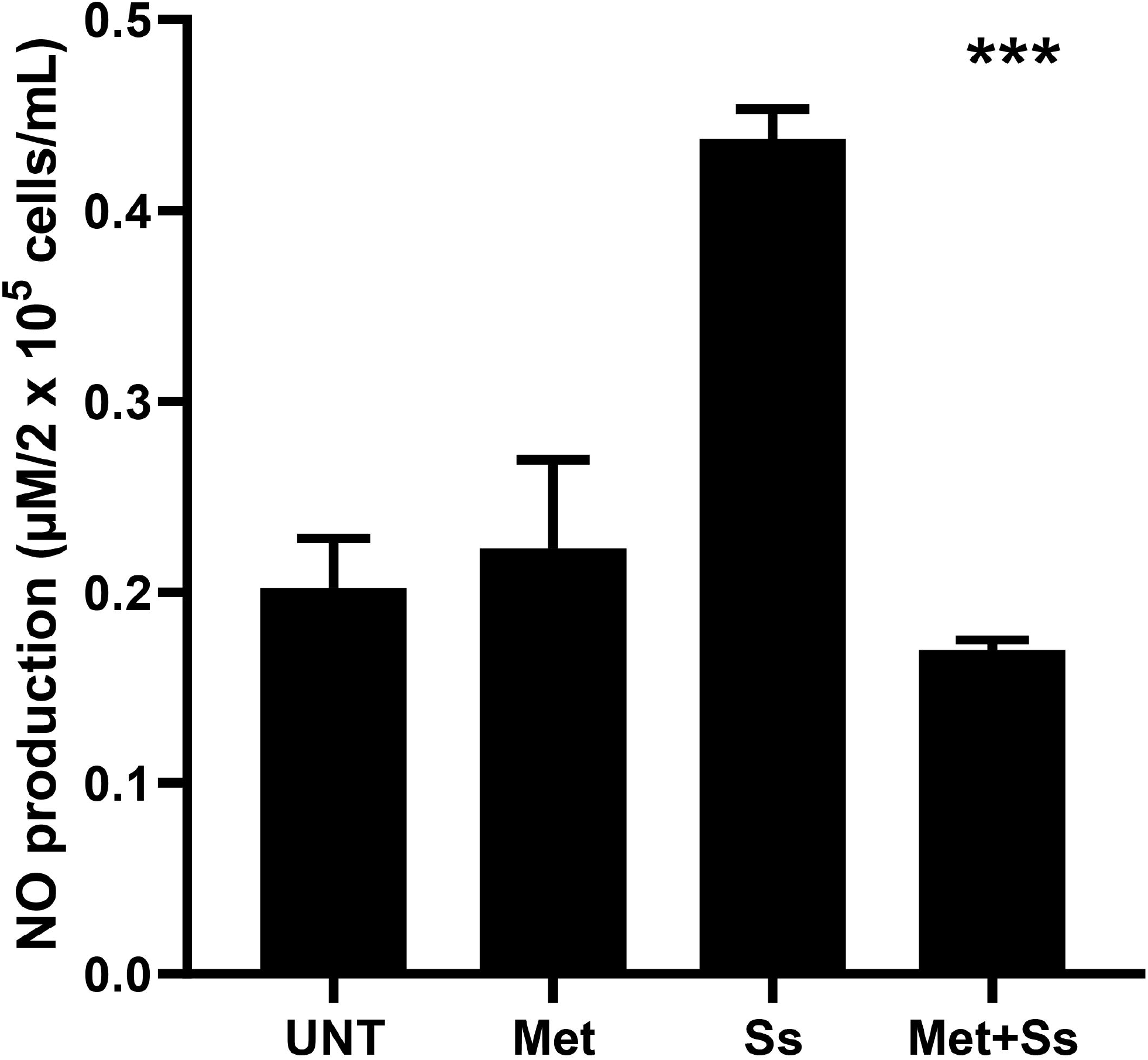
Effect of Met, combined or not to Ss, on the NO production by MOs during T-ALL. NO levels were spectrophotometrically measured by a sensitive Griess method. Asterisks indicate significant differences highlighted between all groups using one-way analysis of variance (ANOVA) test: *\*\*\*P* < 0.001. Data are presented as the mean values with standard error of mean (SEM) of five independent experiments carried out on cell samples (n = 15 in each group). UNT: untreated MO controls, Met: MOs treated with metformin alone, Ss: MOs treated with sodium selenite alone, Met+Ss: MOs treated with metformin combined to sodium selenite, MO: monocyte, NO: nitric oxide, T-ALL: T-cell acute lymphoblastic leukemia.

### 3.2 Met+Ss combination therapy and Ss monotherapy downregulate extracellular calcium ions consumption

As observed in Figure 3, Met treatment induced a slight downregulation of _ec_Ca^2+^ levels *(p* > 0.05), but the difference with Met-untreated MOs reached a significant level when treatment with Met has been combined with Ss *(p* < 0.05). Additionally, MOs treatment with Ss alone induced a highly significant downregulation of _ec_Ca^2+^ levels *(p* < 0.01). Multiple comparisons using one-way ANOVA test gave a *p*-value less than 0.01.

**Figure 3.**
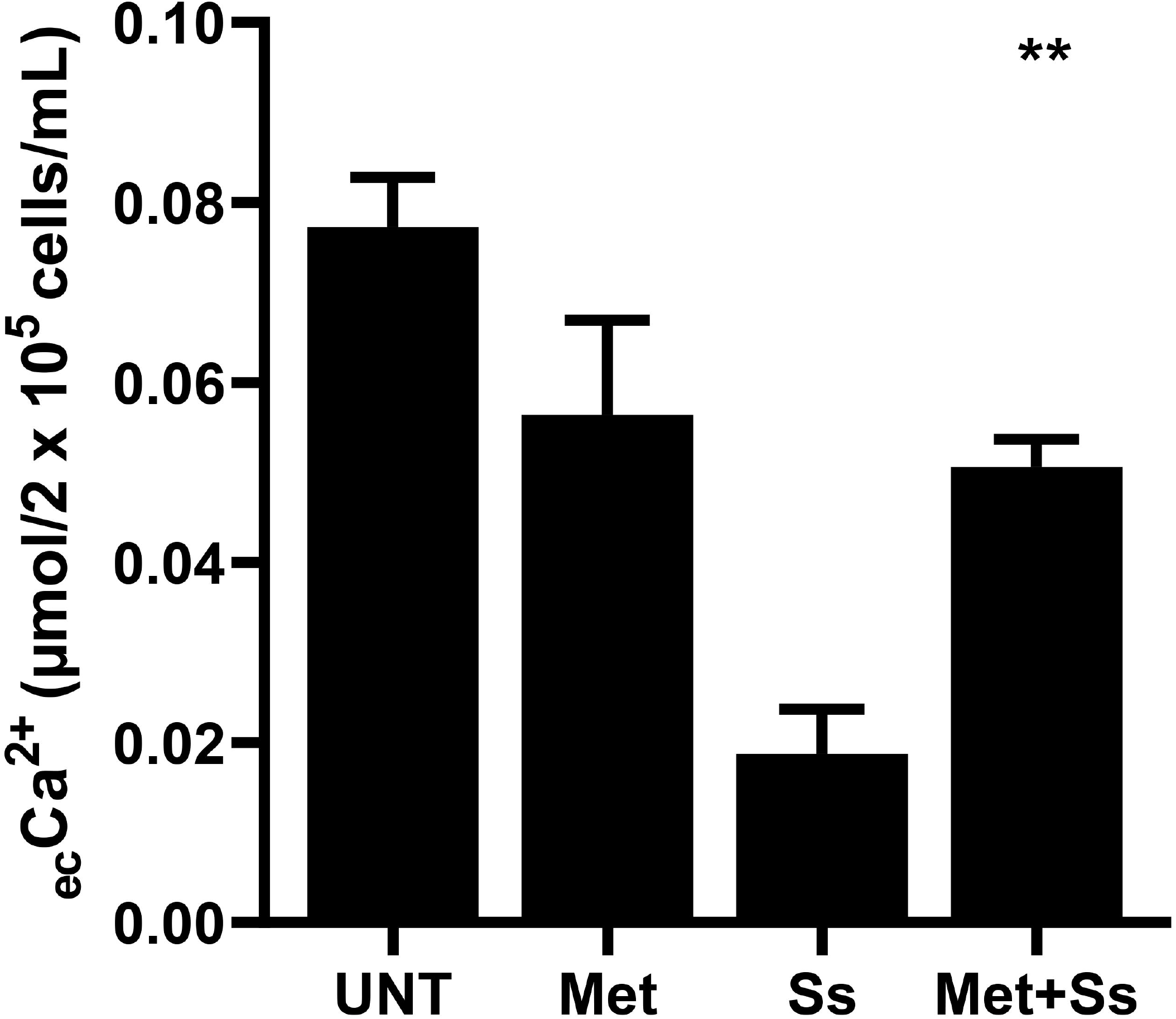
Effect of Met, combined or not to Ss, on the calcium consumption by MOs during T-ALL. MO calcium consumption levels were measured using spectrophotometric method. Asterisks indicate significant differences highlighted between all groups using one-way analysis of variance (ANOVA) test: *\*\*P* < 0.01. Data are presented as the mean values with standard error of mean (SEM) of five independent experiments carried out on cell samples (n = 15 in each group). _ec_Ca^2+^: extracellular calcium ion consumption, UNT: untreated MO controls, Met: MOs treated with metformin alone, Ss: MOs treated with sodium selenite alone, Met+Ss: MOs treated with metformin combined to sodium selenite, MO: monocyte, T-ALL: T-cell acute lymphoblastic leukemia.

### 3.3 Met+Ss combination therapy *versus* Met or Ss monotherapy effects on the relative expression of Bcl-2-associated survival and cytokines production by MOs

As we observe in Figure 4, Met treatment induced a significant upregulation of IL-1β production by T-ALL MOs *(p* < 0.001). Nevertheless, its effect appears to be markedly inverted when combined to Ss treatment (Me+Ss *vs.* Met, *p* < 0.001). Additionally, Ss induced no effect of MO IL-1β levels, either used alone or in supplementation with Met when compared to UNT control MOs *(p* > 0.05). Moreover, neither Met nor Ss treatment has an effect on the relative expression of Bcl-2 *(p* > 0.05). However, combination of Met with Ss induced a significant upregulation of relative Bcl-2 expression in MOs, when comparing with either UNT control MOs or Met-or Ss-treated MOs (*p* < 0.001). Met treatment induced no effect on the production of both IFN-γ and IL-10 by T-ALL MOs. Similarly, Ss has been observed to be without effect on the production of the two cytokines, but when used alone, its induced slight upregulation of IFN-γ production (for all comparisons against control MOs, *p* > 0.05). Additionally, the IL-6 production has changed slightly in Met- and/or Ss-treated MOs, but differences have not reached significant levels when comparing with treated MOs (for all comparisons, *p* > 0.05). Moreover, the levels of TNF-α were significantly upregulated in MOs treated with Met *(p* < 0.05); while, such an effect was reversed by the Met+Ss. Finally, for multiple comparisons using one-ANOVA test, *p*-values were less than 0.001 for IL-1β, less than 0.01 for Bcl-2, greater than 0.05 for IFN-γ, IL-10 and IL-6, and less than 0.05 for TNF-α.

**Figure 4.**
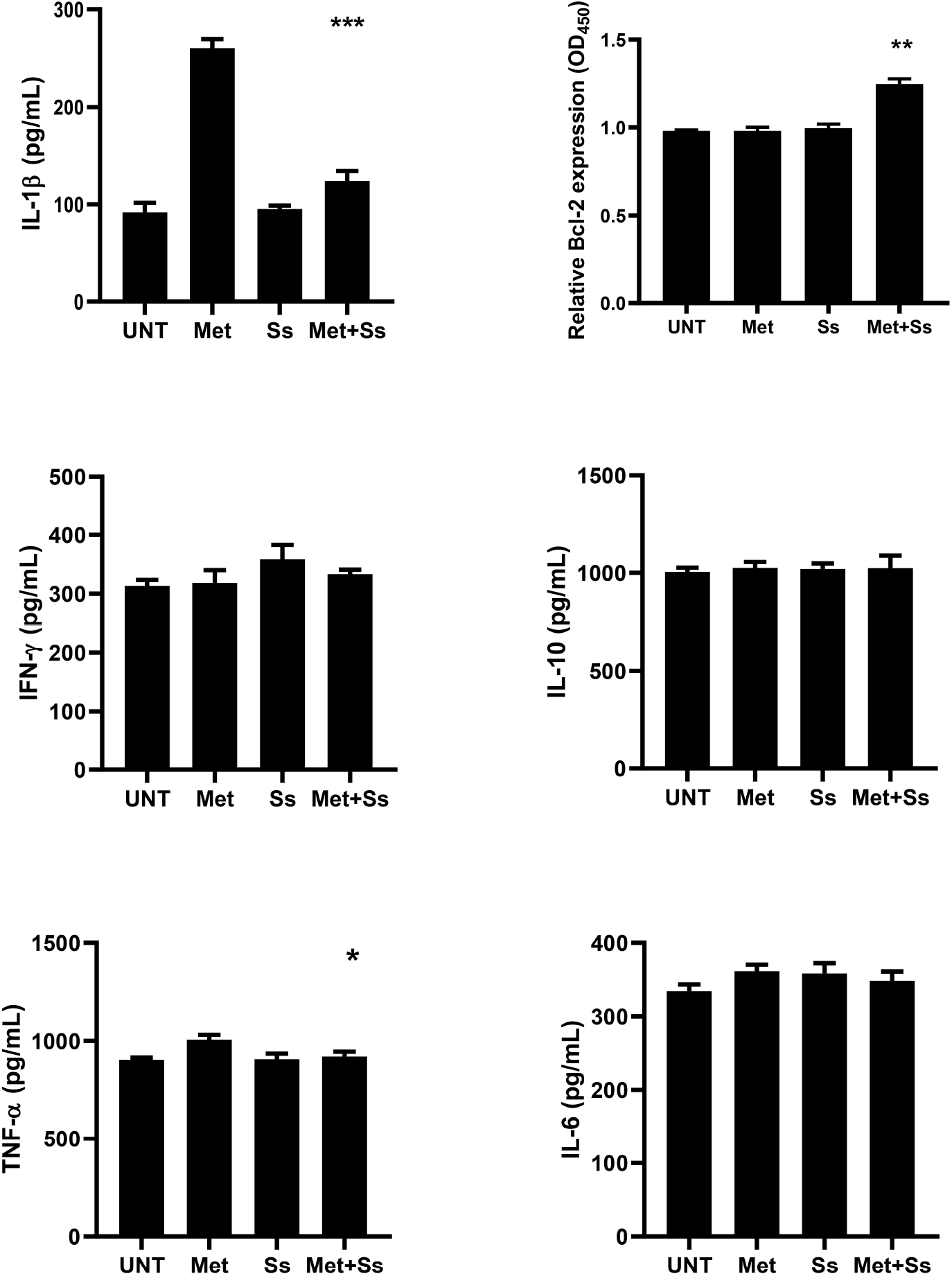
Effect of Met, combined or not to Ss, on the relative expression of Bcl-2-associated survival and cytokines production by MOs during T-ALL. The relative Bcl-2 expression and cytokine levels were measured using enzyme-linked immunosorbent assay (ELISA). Asterisks indicate significant differences highlighted between all groups using one-way analysis of variance (ANOVA) test: *\*P* < 0.05, *\*\*P* < 0.01 and *\*\*\*P* < 0.001. Data are presented as the mean values with standard error of mean (SEM) of three or four independent experiments carried out on cell samples, respectively for the determination of cytokine levels (n = 9 in each group) or the relative Bcl-2 expression (n = 12 in each group). Bcl-2: B cell lymphoma-2, IFN-γ: interferon gamma, IL-10: interleukin 10, IL-1β: interleukin 1 beta, IL-6: interleukin 6, T-ALL: T-cell acute lymphoblastic leukemia, TNF-α: Tumor necrosis factor alpha, UNT: untreated MO controls, Met: MOs treated with metformin alone, Ss: MOs treated with sodium selenite alone, Met+Ss: MOs treated with metformin combined to sodium selenite, MO: monocyte.

## 4. Discussion

Changes in the levels of NO and cytokines derived from circulating monocytes have been demonstrated to have a strong impact on the development and progression of malignant cells. Such changes could be induced by different drugs used alone or in combination. So, combination therapies are often offered for the treatment of the most dreadful diseases, such as cancer, and are frequently first tested on cell cultures [26]. Therefore, this study was focused on the *ex vivo* effect of Met, a biguanide class antitumor drug, combined or not with Ss, on the changes in the production of NO and selected cytokines by circulating human primary MOs of children with T-ALL.

### 4.1 Effect of Met and Ss on the NO production by MOs during T-ALL

NO is involved in the host immune defense system against foreign pathogens and tumor cells [27,28]. In addition to its antimicrobial properties, it can induce apoptosis and cell cycle arrest [29]. It has been reported that its suppressive action on cancer cells could be effective only with high levels, promoting halt cancer growth and acting as a therapeutic agent [30]. Regarding treatment with Met alone, our results corroborate those we recently highlighted about the levels NO production by human granulocyte-macrophage colony-stimulating factor (GM-CSF) MO-derived (GM-MDM) in normal condition [31]. By cons, Ss treatment induced upregulation of NO production when used alone. However, the combination of the two treatments induced a different effect, resulting in a significant downregulation of NO production, suggesting that a combination of the two drugs could affect the expected effects of NO and reduce its effectiveness. Nevertheless, the selenium effect can differ according to the origin of MOs and host condition. Hence, it has previously been reported that selenium supplementation in the form of Ss can induce a downregulation of NO production by mice MO/macrophage-like cell line RAW 264.7 (ATCC TIB-71) [32,33], suggesting that selenium suppresses the nuclear factor that binds to the enhancer element of the immunoglobulin kappa light-chain of activated B-cells (NF-kappa B) activation and interferon regulatory factor 3 (IRF3) induced by toll-like receptor 3 (TLR3) or TLR4 agonists. It would be best to make a kinetics study on Met combined to Ss to highlight the optimal level of NO production when considered as endpoint in malignancy conditions, including T-ALL. Finally, to the best of our knowledge, there are no similar studies to ours, which render difficult to compare our results to others.

### 4.2 Effect of Met and Ss on the consumption of calcium by MOs during T-ALL

Ca^2+^ is universal second messenger, and plays a key role in immune cell signaling [34]. It can modulate both inflammatory immune response, as well as the production of cytokines by MOs-macrophages [35,36]. Our results showed that Met treatment has no marked effect on the calcium ions consumption by MOs from T-ALL condition. Conversely, Ss has been observed to have a significant downregulation effect on the consumption of extracellular calcium ions. Our outcomes suggest that Met and Ss would have an inhibitory effect on the extracellular calcium influx, and consequently on free intracellular calcium that originates from the extracellular medium, which agrees with our recent study carried out on GM-MDM [31]. Therefore, in the process of MO activation during therapy with Met or Ss, other sources of calcium ions, *i.e.,* ergastoplasm and the calcium released from intracellular complexes may be used, which deserves further investigations highlighting the main differences between free and released calcium ions. Additionally, it would be interesting to check whether the low levels in the extracellular calcium consumption is probably due or not to its association with Met, and more particularly, with Ss. Nevertheless, the current results would be of great interest, when we know that the availability of extracellular calcium can inhibit the leukemia cells division, as previously reported [37].

### 4.3 Effect of Met and Ss on the production of IL-1β by MOs during T-ALL

IL-1β is a potent proinflammatory cytokine, which is crucial for the host response against infection and cell damage. It is also considered as the best characterized and the most studied of the 11 members of the IL-1 family [38]. It is produced and secreted by various types of cells, especially MOs and macrophages [39,40]. In our study, we observed that the treatment of MOs with Met alone, or in combination with Ss, induced a marked upregulation of the IL-1β production. Based on previous studies, the effect of Met on the IL-1β production remains still unclear and somewhat paradoxical. The discrepancy between results would be related to the experimental conditions. Therefore, in healthy condition, Met has been demonstrated to downregulate the production of IL-1β in LPS-activated macrophages [41], and, conversely, it has been found without a marked effect in GM-CSF monocyte-derived macrophages; while, Ss has observed to upregulate IL-1β production [8]. Additionally, differences between results should be linked to the type of cells, knowing that human MOs release 22 times more IL-1β per cell than macrophages [40], which corroborate with our results. Moreover, it is well known that, in cancer condition, Met treatment induced a functional phonotypic switching to M1 classically activated, killer macrophage, that is recognized to produce IL-1β [42,43], and attenuated M2 alternatively, activated macrophage [44]. Moreover, a low production of both IL-1β and NO has been observed in tumor-associated macrophage (TAM) cells, which is recognized to promote tumor growth and dissemination in many tumors [45]. Furthermore, it has been reported that an elevated number of MOs occur not only in response to infection, but has also been observed in several chronic inflammatory diseases associated with increased levels of IL-1β. Our results concerning the upregulation of IL-1β production by MOs treated with Met alone, suggest that Met would have a more specific effect on the activation of classical and intermediate MOs, knowing that the classical and intermediate MOs subsets release significantly higher levels of mature IL-1β upon stimulation of LPS compared to the non-classical subset, as reported [46]. Finally, on the basis of the results about Met and Ss combination, we suggest that Ss would have a control action on that of Met.

### 4.4 Effect of Met and Ss on the production of by co-operative ‘immunostimulating/antitumor’ IFN-γ and ‘regulatory’ IL-10 cytokines by MOs during T-ALL

IL-10 is recognized as a pleiotropic cytokine that impact the immune pathology of cancer [47], and as an anti-inflammatory/immunoregulatory cytokine, which is produced by various cells, including MOs [48]. It can act as a rheostat for M1 macrophage glycolytic commitment [49], and inhibit the production of proinflammatory cytokines, such as IFN-γ, TNF-α and IL-1β [50]. Nevertheless, there are some reports about its co-operative action with ‘immunostimulatory’ cytokines [51], although recent evidence suggesting requirement of IL-10 for the IFN-γ anti-tumoral action, as well as for T-helper cell functions, T-cell immune surveillance, and suppression of cancer-associated inflammation [20]. That leads to open new avenues of research on the dual role of IL-10.

Although involvement of IFN-γ in tumor immune surveillance as antiproliferative and pro-apoptotic cytokine, clinical trials remain with limited success regarding its effects on progression-free survival. This could be due to the fact that it is not fully with anti-tumor effects, and that it could play an adverse role. Indeed, it has been suggested that it might also have a protumorigenic role, which results from changes in the activation of its signaling pathways [52,53]. In the current study, Met treatment, combined or not with Ss, induced no change in the production of IFN-γ by MOs. So it has been reported that Met treatment, not only affects the differentiation and maturation of MOs into macrophages, but also attenuates the production of proinflammatory cytokines [54,55]. Of note, it would be of great interest to check whether or not the Met effects are likely to be corrupted or affected by inhibitory signals during T-ALL.

Our results show similarity with others regarding Met effect on the IL-10 release in patients with tuberculosis or type 2 diabetes [56], and discordance with that of a study carried out on macrophages from obese diabetics [57,58]. On the other hand, treatment combination approaches have been explored to boost innate immune activation. In this study, however, there were no significant changes in both IFN-γ and IL-10 production by MOs from T-ALL patients when combining Met to Ss. These results open new perspectives investigating whether the production of IL-10 by MO during T-ALL depends on that of IFN-γ. If so, new evidence on the immunomodulatory effects of Met will be provided.

### 4.5 Effect of Met and Ss treatment on the production of IL-6 by MOs during T-ALL

IL-6, a proinflammatory cytokine with pleiotropic biological properties, participates in several physiological and immune functions, including regulation of activities of immune cells and hematopoiesis [59]. Additionally, IL-6 can be produced in response to local proinflammatory cytokines such as TNF-α [60]. Moreover, IL-6 signaling in the tumor microenvironment sustaining angiogenesis and tumor evasion of immune surveillance, supporting cancer cell proliferation, survival, and metastatic dissemination. In marked contrast with the evidence that IL-6 promotes cancer cell development, it has been reported that it can also play an antitumoral role, and orchestrate effective anti-tumor immunity in tumor microenvironment [61], by resculpting the T-cell immune response [62]. Additionally, IL-6 signaling in macrophages has recently been reported to be critical for tumor tissue destruction under circumstances of immunotherapy [63]. So in our study, the induction of upregulation (albeit slightly) of monocytic IL-6 by Met, used alone or in combination with Ss, should open up new avenues of research to know if this would have a beneficial or, on the contrary, deleterious effect. These results should also lead to check whether there is or not a repercussion on the MO maturation into macrophages, and the polarization of these ones, knowing that IL-6 has been shown to be a cytokine that controls macrophage alternative activation [64].

### 4.6 Effect of Met and Ss treatment on the production of the pro- and antitumoral TNF-α by MOs during T-ALL

In the past, it has been believed that TNF-α act only as a tumoricidal cytokine [65,66]. Numerous studies have shown, however, that TNF-α may have a dual role with pro- or anti-tumorigenic effect, and can also promote carcinogenesis in most cancer cells that are usually resistant to TNF-α-induced cytotoxicity [67]. Thus, it has also been suggested that blocking TNF-α in combination with immunotherapy would be one of the therapeutic strategies for improving the antitumor effect and preventing or treating adverse effects linked to the immune system. For its antitumor effect, TNFα antitumor mechanisms are varied, including: (i) mediating cell apoptosis, (ii) directing TAMs towards the M1 antitumor phenotype, (iii) guiding neutrophils and monocytes to tumor sites, (iv) activating macrophages and inhibiting the differentiation of MOs into immunosuppressive phenotypes, (v) and inducing disruption of tumor vascularization [68]. In the present study, we observed that Met treatment induced a marked upregulation of TNF-α production by MOs from T-ALL; while, neither Ss treatment alone nor the Met+Ss combination induced a change in the levels of TNF-α production. Paradoxically, it has previously been reported that Met attenuated the production of TNF-α by macrophages, as well as in inflammatory condition, like obesity [69]. Nevertheless, to the best of our knowledge, there are no similar studies on MOs from T-ALL allowing a perfect comparison with our results.

### 4.7 Effect of Met and Ss on the expression of the anti-apoptotic Bcl-2 protein by MOs during T-ALL

The Bcl-2 transcription factor plays an important role in promoting cell survival and in inhibiting the actions of pro-apoptotic proteins [70]. Mechanistically, Bcl-2 prevents oligomerization of Bax/Bak, which leads to the release of a number of apoptogenic mitochondrial molecules [71]. Bcl-2 has recently been shown to improve the persistence of chimeric antigen receptor-modified T-cells (CAR-T) by reducing apoptosis induced by activation in a mouse xenograft lymphoma model, thereby improved anti-tumor activity [72]. In our study, we observed that neither Met, nor Ss affected the levels of MO Bcl-2 expression; whereas, Met+Ss combined treatment induced increased Bcl-2 expression. When we see the results of IL-1β, we might suggest that Met+Ss combination potentiated the physiological mechanism of Bcl-2, which is known to reduce the activity of caspase-1, and therefore, the IL-1β production [73]. Additionally, the downregulated levels of IL-1β we reported above would be mainly related to the effect of Ss, which has recently been demonstrated with immunomodulatory effects on Met in macrophage in healthy condition [8]. Nevertheless, increasing the level of Bcl-2 expression, along with decreasing IL-1β production requires a synergistic effect combining Met and Ss. Therefore, it would be wise to test such a combination with different doses of both Met and Ss.

## 5. Conclusions and Future Prospects

T-ALL results from malignant transformation of T-cell progenitors, affecting both childhood and adult. Poor prognosis has been linked to leukemic T-cells infiltration in organs after their transendothelial migration. This can promote their interaction with various immune cells, including MOs. The latest might have dual roles leading to either cancer cell lysis or development, depending to cell microenvironment. So enhancing their functional activities, including production of NO and antitumor cytokines would be of great interest, especially in the context of a tumor-induced immunoediting, when both MOs and macrophages can undergo a phenotypic switch into immunosuppressive cells. Herein, we demonstrated that although Met or Ss-treated MOs has without notable effect on functional phenotypic activities of human MOs regarding pro- or anti-tumor cytokines production during T-ALL, Met combination with Ss appears to have a marked effect on MOs survival and downregulation of IL-1β production. Finally, the Met action would be reversed by Ss treatment, especially on the release of both IL-1β and NO. *In fine,* this study should pave the way for other *ex vivo* or *in vivo* studies exploring the therapeutic effects of Met and its combination with Ss during the interaction of leukemic T-cell with different human MO subtypes during T-ALL, *i.e.,* the so-called classical inflammatory (moM1, CD14^++^/CD16^-^), intermediate (moM2, CD14^++^/CD16^+^) and patrolling non-classical MOs (moM3, CD14^+^/CD16^++^). In addition, our outcomes regarding the downregulation effect on the _ec_Ca^2+^ in Met+Ss- or Ss-treated MOs deserve investigation checking if Met or Met+Ss action on calcium signaling depends or not on calcium release-activated channels (CRAC).

## Acknowledgments

The authors would like to express their deep thanks to the parents of participants for their invaluable patience. They are also grateful to the Thematic Agency of Research in Health Sciences for their financial support.

## Competing Interests

The authors have declared that no competing interests exist.

---

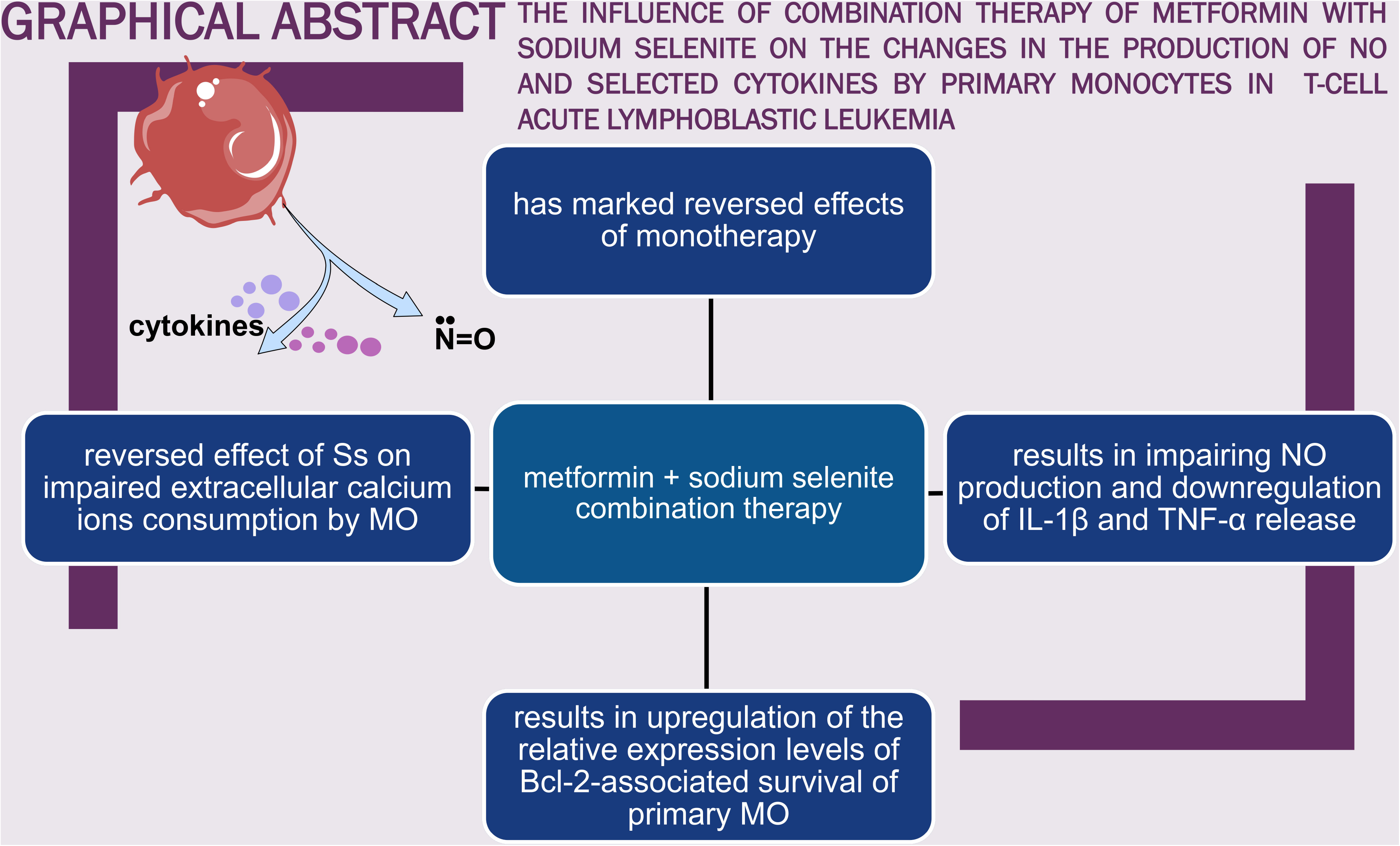

## Notes

### Competing Interest Statement

The authors have declared no competing interest.

